# Biomolecular Condensates can Induce Local Membrane Potentials

**DOI:** 10.1101/2024.12.27.630407

**Authors:** Anthony Gurunian, Keren Lasker, Ashok A. Deniz

## Abstract

Biomolecular condensates are a ubiquitous component of cells, known for their ability to selectively partition and compartmentalize biomolecules without the need for a lipid membrane. Nevertheless, condensates have been shown to interact with lipid membranes in diverse biological processes, such as autophagy and T-cell activation. Since many condensates are known to have a net surface charge density and associated electric potential(s), we hypothesized that they can induce a local membrane potential. Using an electrochromic dye, we demonstrate that poly-lysine/ATP condensates induce a localized membrane potential in Giant Unilamellar Vesicles. This effect diminishes with increasing salt concentration and higher ATP-to-poly-lysine ratios, underscoring the key role of condensate charge. Numerical modeling of the condensate-membrane interface using an electro-thermodynamic framework supports our experimental findings and highlights parameters expected to play a key role in the effect. These results have broad implications for biological processes regulated by membrane potential, particularly in contexts such as neuronal signaling, where condensate interactions with membranes may play a previously unrecognized regulatory role.

## Introduction

In the last decade, biomolecular condensates have been recognized as a ubiquitous feature of cells with roles in many cellular functions ranging from nucleolar function [1, 2], transcription and signaling [3], to nuclear pore permeability [4]. Biological condensation has also been linked to cell misfunction and disease [5], for example through altered condensate properties [2, 6, 7], protein misfolding in condensates [7-12], and viral condensates [13, 14]. One major focus in functional studies of condensates has been their *partitioning property* (the ability to selectively concentrate or exclude certain molecules). Partitioning of biomolecules has the effect of spatially organizing cellular processes, and has been shown to increase the rate of biomolecular reactions in some cases [15]. There have also been substantial efforts in some other directions, such as condensate material, interfacial and mechanical properties, with important links to biological function and disease being uncovered [1, 2, 6, 16-19]. Much less is known about other properties of condensates and how they relate to biological functions.

Recently, the electrical properties of condensates have gained some attention. Electrophoretic light scattering has been used to measure condensate zeta potentials with values in the range of -42 mV to +42 mV [20-22]. Microelectrophoresis was used to measure polylysine/polyaspartic acid condensate zeta potentials with a magnitude of 2.5 mV or 20 – 80 mV depending on which model was used to convert electrophoretic velocity to zeta potential [23]. Theoretical work has shown that the interior of a condensate is charge-neutral, while the condensate surface has a net charge resulting in interfacial potentials and/or Donnan potentials on the order of 1-10 k_B_T/e [24], which is 25-250 mV at 25°C. Simultaneously, several papers have investigated interactions between membranes and condensates including membrane wetting and endocytosis [20], membrane tubulation [25], and transmembrane coupling of condensates [26]. In light of these observations, we hypothesized that the wetting of membranes by condensates might alter the local membrane potential.

In classical electrophysiology, the (intra)membrane potential (V_intramembrane_) is usually assumed to be equivalent to the transmembrane potential (V_TM_) which arises due to asymmetric ion concentrations following the Goldman-Hodgkin-Katz (GHK) equation [27]. However, intramembrane potentials can also arise as a result of asymmetric surface potentials (V_surface_) or asymmetric dipole potentials (V_dipole_) [28-30]. The surface potential is primarily due to charged lipids such as phosphatidic acid (PA), while the dipole potential is thought to arise from oriented water molecules [31]. If the wetting of a membrane by a condensate alters the surface potential and/or dipole potential on one side of the membrane, it will result in a *local* intramembrane potential (Figure 1).

**Figure 1:**
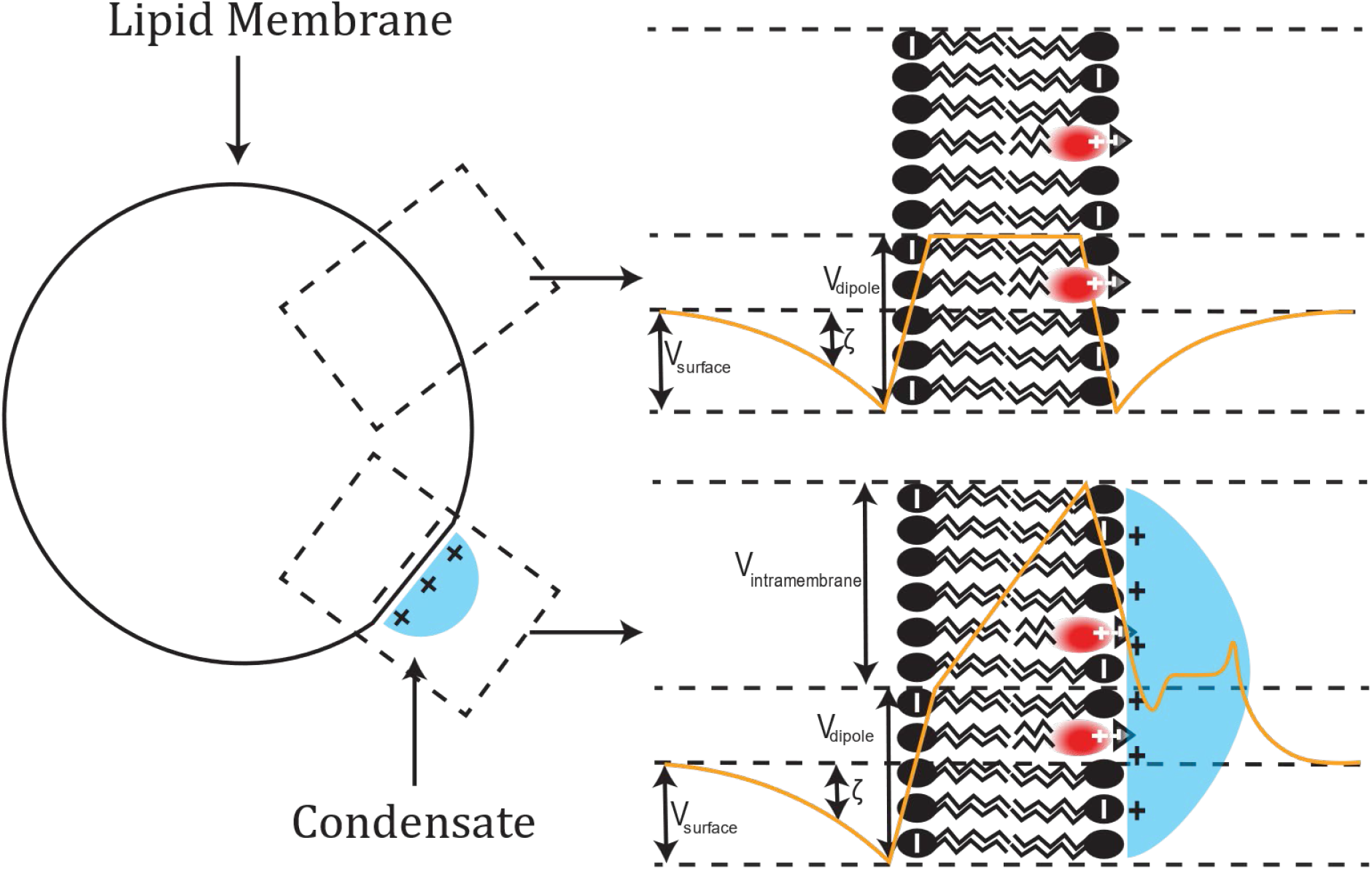
Schematic illustration of hypothetical electric potential profile (orange) informed by numerical simulations, across the lipid membrane (black), with associated condensate (blue), and di-8-anepps (red). **V**_**intramembrane**_ – intramembrane potential, **V**_**surface**_ – surface potential, **V**_**dipole**_ – dipole potential, **ζ** – zeta potential

Previously, it has been shown that when basic peptides bind to membranes, they can induce domains enriched in acidic lipids at certain concentration regimes [32]. The domains enriched in acidic lipids were theoretically predicted to have a higher surface potential [32]. Binding of poly-lysine to negatively charged lipid membranes has been shown to reverse the zeta potential of lipid vesicles and alter the membrane dipole potential of planar lipid bilayers, in a concentration dependent manner [33]. In analogy to the binding of basic polypeptides to membranes [32, 33], a local difference in surface potential could arise simply due to the much higher polyelectrolyte concentration in the condensate compared to the dilute phase. A recent paper used LAURDAN, a solvatochromic dye, to show condensate induced local changes in membrane *hydration* [34]. Changes in the average orientation, concentration, or rotational dynamics of water molecules at the membrane-condensate interface could also result in an altered dipole potential. Finally, the existence of a condensate Donnan potential [24, 35], creates different local bulk phase electric potential, which could result in a transmembrane potential.

Recently, di-4-anepps was used to show *global* changes in membrane potential in E. coli due to global changes in intracellular ion concentration [36]. However, since ions are free to diffuse in all directions, it should be impossible to have any kind of *local* difference in membrane potential at equilibrium within the framework of the GHK equation. Here, we use di-8-anepps, an electrochromic dye, to show that condensates can alter the *local* membrane potential. This could have large implications for the regulation of voltage gated ion channels and other electric field sensitive membrane proteins, which we elaborate on in the discussion.

## Results

### Condensates can induce local membrane potentials

To evaluate whether condensates can induce membrane potentials, we started with a model system consisting of GUVs (Giant Unilamellar Vesicles) and polyK/ATP complex coacervates. We stained the GUV membranes with an electrochromic dye, di-8-anepps [37], which shifts its excitation and emission spectra in the presence of a local electric field. The shift in absorption frequency can be described by the equation:

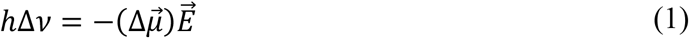

where *h* is Planck’s constant, 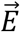 is the local electric field, and 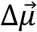 is the change in dipole moment between the ground state and the excited state [38, 39]. Following previous work [37], we use the excitation ratio R_440/514_ to quantify the shift in the excitation spectrum. Compared to absolute intensity measurements, the ratiometric approach has the advantage of being independent of local variations in dye concentration. Also, we chose di-8-anepps because of its very low rate of flipping between leaflets of the lipid bilayer [40]. In the following experiments, we assume that most of the dye is retained in the outer leaflet during the experiment.

By mixing GUVs composed of zwitterionic POPC and anionic PA, with positively charged poly-lysine/ATP complex coacervates, we observed membrane wetting and changes in curvature (Figure 2A-C). We also observed a striking local red shift of the dye excitation ratio R_440/514_, indicating a local change in membrane potential (Figure 2D-F). According to our model (Figure 1), the adsorption of the positively charged condensate outside the GUV creates an electric field directed toward the inside of the GUV. Since the dye is oriented with the sulfonate group pointing outward, the electric field vector is parallel to the vector 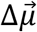 [39], resulting in a negative change in frequency, or red shift, according to equation 1. Note that alternatively, if the condensate has the effect of lowering the membrane dipole potential (V_dipole_), this would also result in a red shift since the electric field due to the dipole potential is directed antiparallel to the vector 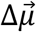.

**Figure 2:**
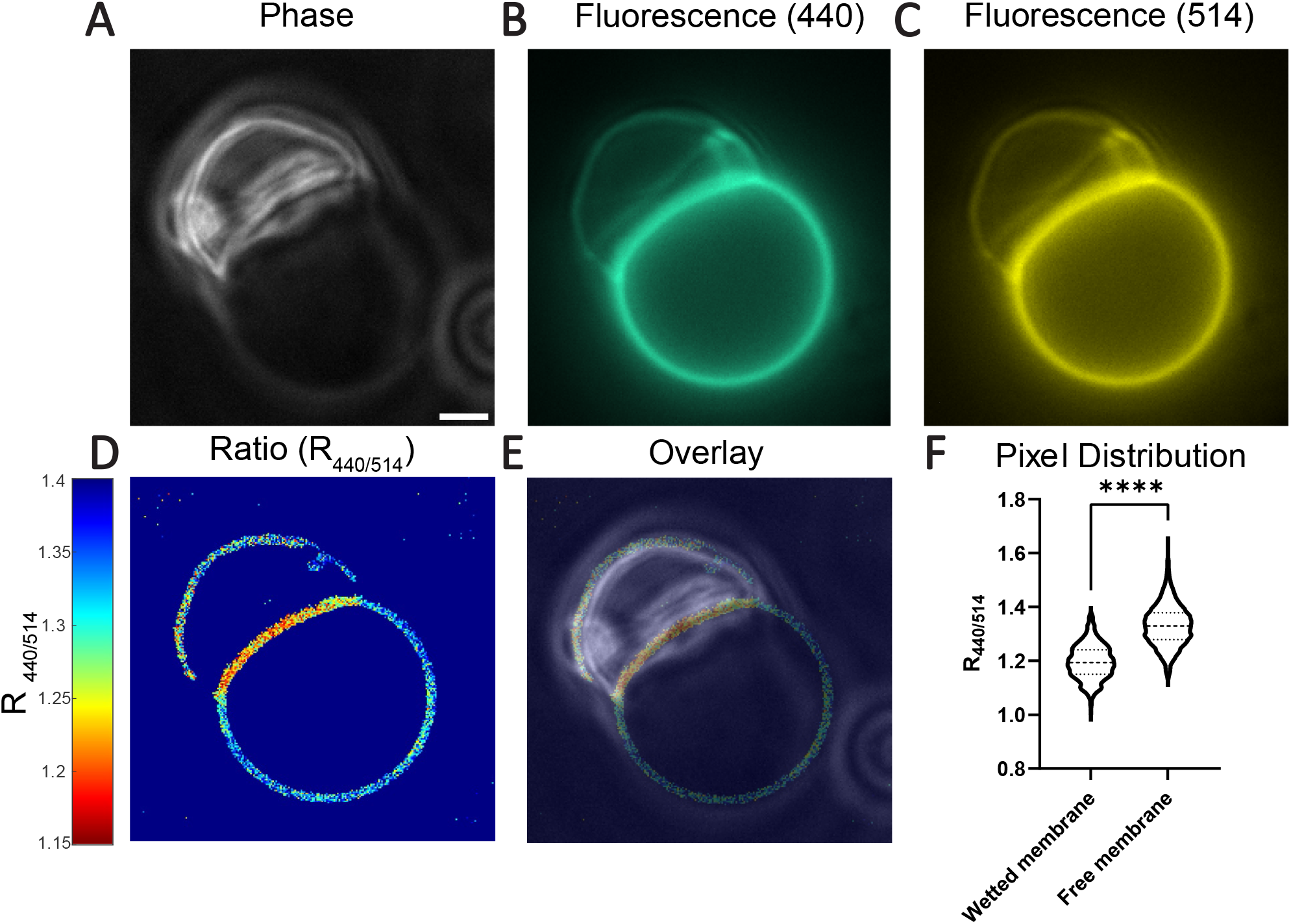
Example of wetting of GUVs composed of 80% POPC, 20% PA by polyK/ATP condensates (20 µM polyK-50, 20 µM polyK-100, 1 mM ATP, 2.5 mM MgCl_2_). (A) Phase contrast image of condensate wetting GUV. (B,C) Fluorescence images of GUV membrane labeled by di-8-anepps, with 440 nm and 514 nm excitation respectively. (D) Thresholded raw ratiometric image, (E) Overlay of phase contrast image and ratiometric image. (F) Distribution of background subtracted pixel intensity ratios in wetted membrane segment and free membrane segment (unpaired t-test with Welch’s correction, p < 0.0001,^****^), Scale Bar 2 µm.

### The induced effects are directly related to condensate charge, along with other contributions

To demonstrate that our observed effect is directly related to the charge of the condensate, we first titrated the concentration of ATP. Increasing the concentration of ATP should decrease the magnitude of the condensate surface charge density, by changing the polyK/ATP ratio [23]. Figure 3 (A, B, C, E) shows that the less positively charged coacervates wet the membrane to a lesser extent, and induce a lower membrane potential. Next, we performed another control, where we increased salt concentration, which should screen the condensate interactions with the membrane, and lower the condensate Donnan and interfacial potentials [24]. Figure 3 (D, F) shows exactly this trend. These control experiments strongly support the (expected) major role of condensate charge on the induced local membrane potentials.

**Figure 3:**
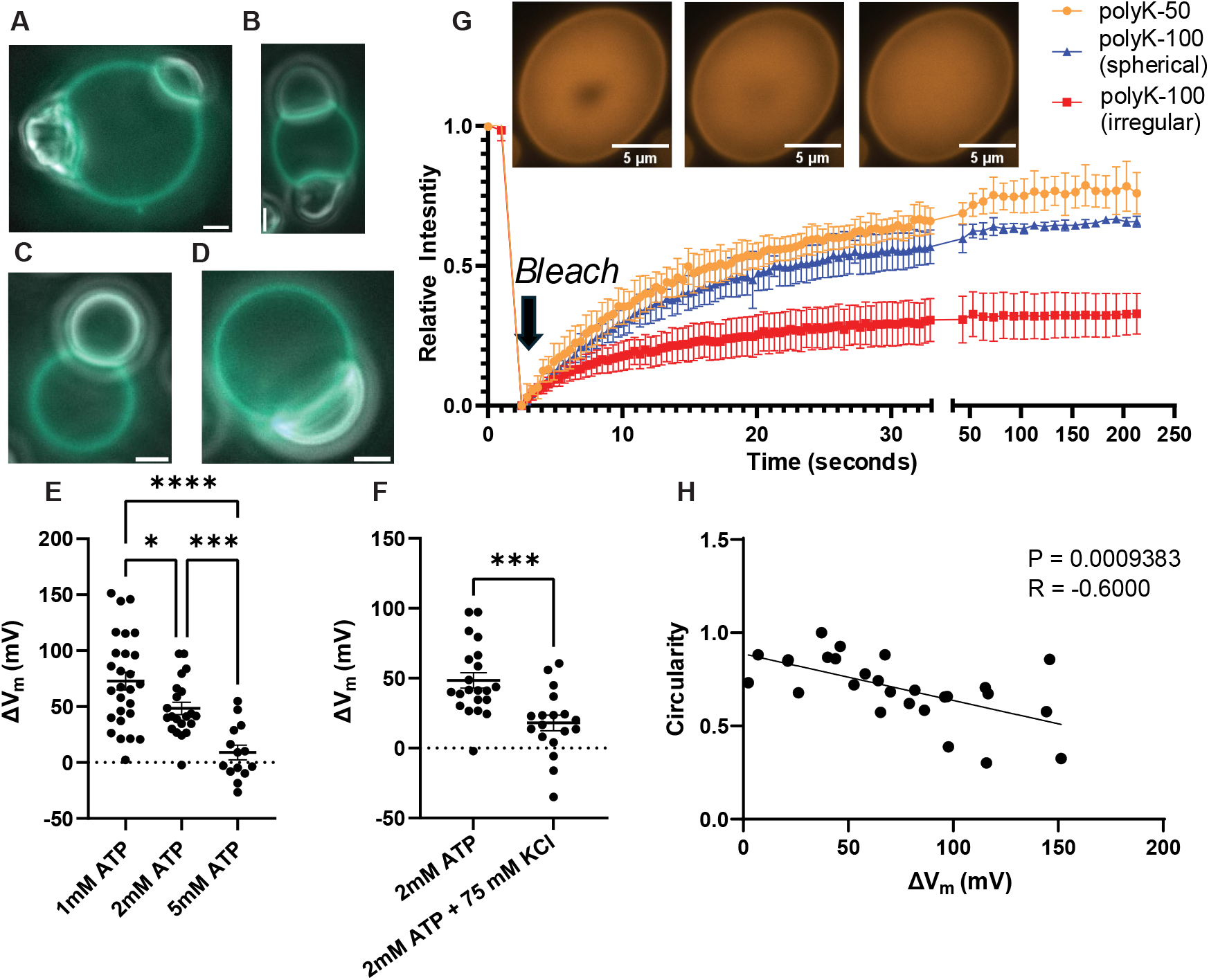
GUVs are composed of 80% POPC, 20% PA, and polyK/ATP condensates are formed with 20 µM polyK-50, 20 µM polyK-100, x mM ATP, 2.5 mM MgCl_2_. (A) 1 mM ATP (B) 2 mM ATP (C) 5 mM ATP (D) 2 mM ATP + 75 mM KCL (E) Summary figure of ATP titration (One-way ANOVA and Dunnett’s T3 multiple comparisons test (p<0.0001,^****^ | p<0.001,^***^ | p<0.05,*). (F) Effect of Salt (unpaired t-test with Welch’s correction, p < 0.001,^***^) (G) FRAP curves of condensates corresponding to 30 µM polyK-100 1.275 µM ATP, 2.5 mM MgCl_2_ or 60 µM polyK-50 1.275 µM ATP, 2.5 mM MgCl_2_. Inset images correspond to polyK-50 condition. (H) Correlation of condensate circularity with ΔV_m_. Condensates (20 µM polyK-50, 20 µM polyK-100, 1 mM ATP, 2.5 mM MgCl_2_). R value is Pearson’s correlation coefficient. Scale Bar 2 µm. Error bars represent SEM.

We wondered what other characteristics of condensates might affect the magnitude of the change in membrane potential (Δ*V*_*m*_), and whether they may be responsible for the large standard deviations in our measurements of membrane potential (Fig. 3 e,f). Throughout the course of our experiments, we noticed that sometimes within the same condition (e.g., Fig. 3 a,b) there were coexisting populations of spherical, liquid-like condensates, and irregularly shaped, gel-like condensates. We confirmed that irregularly shaped condensates had a higher viscosity using FRAP (Fluorescent recovery after photobleaching). We then measured the condensate geometric parameters (circularity, solidity, roundness, aspect ratio; see Figure S2 for definitions) and found moderate correlations with Δ*V*_*m*_ (Figure 3h, S2); the correlation with circularity (4π×Area/perimeter^2^; Figure 3h) was particularly good (r = -0.6, p < 0.001), suggesting that both shape elongation and irregularities of the shape boundary are important. Since the shape of a condensate changes when it wets a lipid membrane, with less round shapes and higher aspect ratios typically reflecting greater wetting and stronger interactions [20], some of our observed correlations may be simply because condensates with a greater charge have a stronger interaction with the membrane.

Together, our experimental results demonstrate the induction of local membrane potentials by interacting condensates. Our results also directly show the key role of condensate charge in this effect, with other condensate parameters having an influence.

### Analytical framework and simulations to explore qualitative features of the induced potentials

Finally, to complement our experimental work, we developed an analytical theory framework and used it to carry out numerical calculations to determine the theoretical electrical potential profile at the membrane-condensate interface. We build off an existing model which couples the Poisson equation with a free energy functional combining the Flory-Huggins free energy, with electrostatic free energy and concentration gradient terms [24], and ask what happens in the presence of a charged membrane? Figure 4a shows a schematic illustration of our model geometry and results. The adsorption of positively charged polycations to the negatively charged membrane results in a charge reversal at the membrane and a large positive surface potential at the condensate-membrane interface (Figure 4d, x = 0), while the electric potential at the membrane adjacent to the dilute phase (Figure 4d, x = 40) is much lower. Simultaneously negative charge accumulates immediately adjacent to the membrane, resulting in a net negative charge density (Figure 4c), and a large drop in the electric potential (Figure 4d, x = 2). For the case where the Flory-Huggins interaction parameter is *χ*_12_ = −5, −7, with a more negative value of *χ*_12_ indicating a stronger interaction between the coacervating components, there is also an interfacial component to the electric potential between the dense and dilute phases which has been observed previously [24]. Together, this results in a polarization of the condensate, and a non-negligible electric field 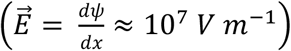 within the bulk of the condensate. This could have implications for the conformation of client proteins within the condensate, which would tend to align their charged groups with the electric field. However, we emphasize that our simulation corresponds to a condensate on the length scale of tens of nanometers, and the effect may be different for larger condensates. Overall, our model and simulations provide additional insight into the nanoscale profile of the electric field at the condensate-membrane interface and makes the prediction that condensates with a stronger interaction parameter *χ*_12_ will result in a larger difference in membrane potential (Figure 4D).

**Figure 4:**
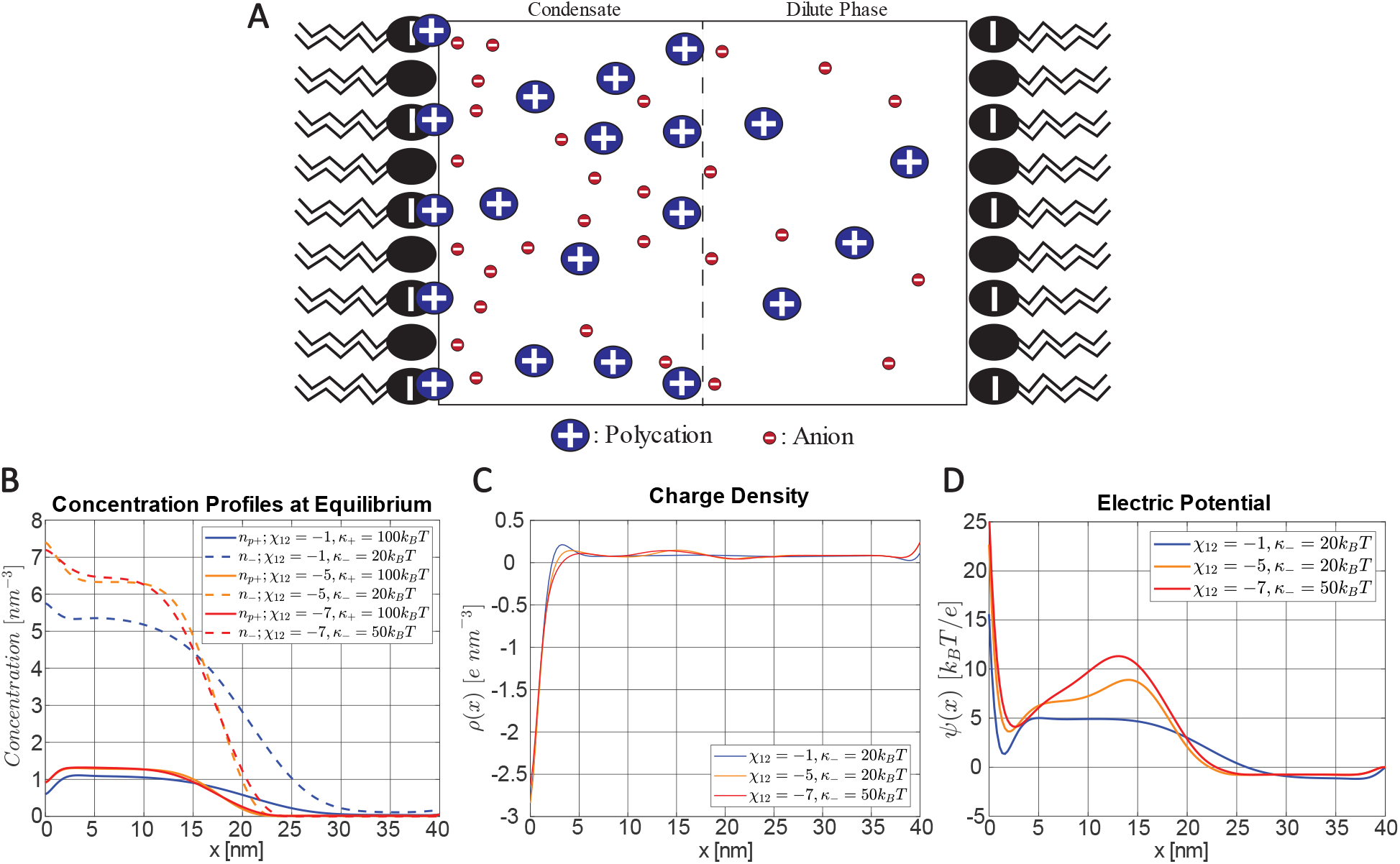
Numerical calculations of a condensate in the presence of charged membranes. (p+) indicates the polycation while (-) indicates the anion. Parameters used for calculation: *κ*_*p*+_ = 100*k*_*B*_*T*; *κ*_−_ = 20*k*_*B*_*T*; *χ*_13_ = 1.5; *χ*_23_ = −0.09; *N*1 = 20; *N*2 = 1; *N*3 = 1; *v* = 0.03 *nm*^3^; *z*1 = 5; *z*2 = −1; *z*3 = 0; *A*_*L*_ = 0.5 *nm*^2^; *f* = 0.5; *K*_*p*+_ = 5. **a**. Equilibrium concentration profiles. Region with high concentrations corresponds to the dense phase (condensate), region with low concentrations corresponds to the dilute phase. **b**. Equilibrium charge density. Separation of charges leads to electric potential. **c**. Equilibrium Electric Potential.

## Discussion

We have shown for the first time that it is possible to generate local differences in membrane potentials at equilibrium. Traditionally, the cell has often been assumed to be analogous to a membrane bound vesicle containing a dilute solution of ions, proteins, and nucleic acids. Since accumulating ions in localized regions is highly disfavored due to the large entropic cost, the concept of local membrane potential has been largely neglected, except for the well-known case of the action potential, which is a transient. Condensates, due to their electrical properties (Donnan potential, Interfacial potential), and the property of being spatially segregated units, are able to induce local membrane potentials where they come into contact with, and wet, lipid membranes. We emphasize that unilateral changes in the dipole potential and surface potential are also able to induce an intramembrane potential, and in future work we will seek to determine which component electric potential(s) are responsible for our observations. Note that the dipole potential, since it occurs over such a short distance, has a larger electric field associated with it (E = V/d), so di-8-anepps is particularly sensitive to the dipole potential.

Previous observations of local variations of membrane potential were all in cells and have been typically attributed to local variations in dipole potential and lipid composition [41, 42]. Also, we should note that others have recently hinted at the converse of our observation, that variations in the dipole potential may drive condensate association with certain membrane regions [43]. We considered the possibility of lipid demixing and accumulation of phosphatidic acid (PA) as an alternative explanation for our results, but PA was previously shown to increase the dipole potential and R_420/520_ [44, 45], whereas we see the opposite at the wetted membrane region. Also, we have seen no indications of lipid demixing in similar systems with fluorescently labeled lipids [20]. Our observations are also not due to a condensate induced increase in lipid packing [34], since this would also cause an opposite change in R_420/520_. [46]. Finally, we did a control experiment replacing the zwitterionic lipid POPC with DOPC (data not shown), and were still able to recapitulate our results.

Ermakov and coworkers have extensively studied the interaction of poly-lysines with lipid membranes [33, 47]. Using zeta potential measurements of liposomes, they observed a sudden, sharp increase in zeta potential at a lysine (monomer) concentration of approximately 200 µM, which corresponds to 2 µM polyK-100. In terms of our results, if the condensate (dense phase) concentration is higher than this cutoff, and the dilute phase concentration is less than the cutoff, then our observed differences in membrane potential could be simply explained by differences in surface potential. While our model suggests poly-lysine also adsorbs to the membrane adjacent to the dilute phase (Figure 4), we see no evidence (within our experimental sensitivity) of poly-lysine adsorption to free membrane regions using condensates doped with labeled polyK-Atto565 (Figure S3), and in similar systems used by others [20]. Importantly, polyK-Atto565 by itself does adsorb to GUVs in the absence of ATP (Figure S3).

In addition, using the intramembranous field compensation (IFC) method with planar lipid bilayers, Ermakov and coworkers observed that changes in boundary potential upon titration with lysine were right shifted compared to changes in zeta potential, and attributed this to opposite changes in the dipole potential [33, 47]. Using molecular dynamics simulations, they showed that lysine molecules displace water molecules which previously made hydrogen bonds with lipid phosphate groups, and since water molecules are thought to be responsible for the dipole potential [31], it was concluded that changes in the number and/or orientation of water molecules is responsible for the decrease in dipole potential [33, 47].

Local changes in membrane potential could have large implications for the function of ion channels and other transmembrane proteins which are regulated by voltage. For example, the class of voltage-gated ion channels change their conformation depending on the intramembrane potential and consequently control the flux of ions in and out of the cell. Therefore, local differences in membrane potential could facilitate localized flux of ions, permitting precise spatiotemporal control over cell signaling (Figure 5). On a different level, the basic equations for the flux of ions should be modified to account for the presence of biomolecular condensates. The flux of ions has traditionally been given as the sum of the diffusive flux determined by the concentration gradient, and the electrophoretic flux determined by the voltage gradient [27]. In the presence of condensates, this should be modified to include a chemical potential gradient 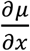 due to the differential partitioning of specific ions in condensates [35, 48], and a modified voltage gradient 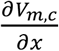 taking into account the condensate induced membrane potential. We can express the modified flux as,

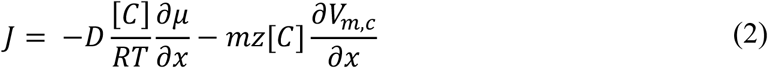

where D is the diffusion coefficient, [*C*] is the ion concentration, R is the gas constant, T is the temperature, m is the ion mobility, and z is the ion valence. One interesting example of a condensate associated with ion channels is the RIM/RIM-BP condensate which clusters voltage-gated calcium channels (VGCCs) at the neuronal pre-synapse [49]. Future work can examine whether this condensate can regulate VGCCs by inducing a membrane potential.

**Figure 5:**
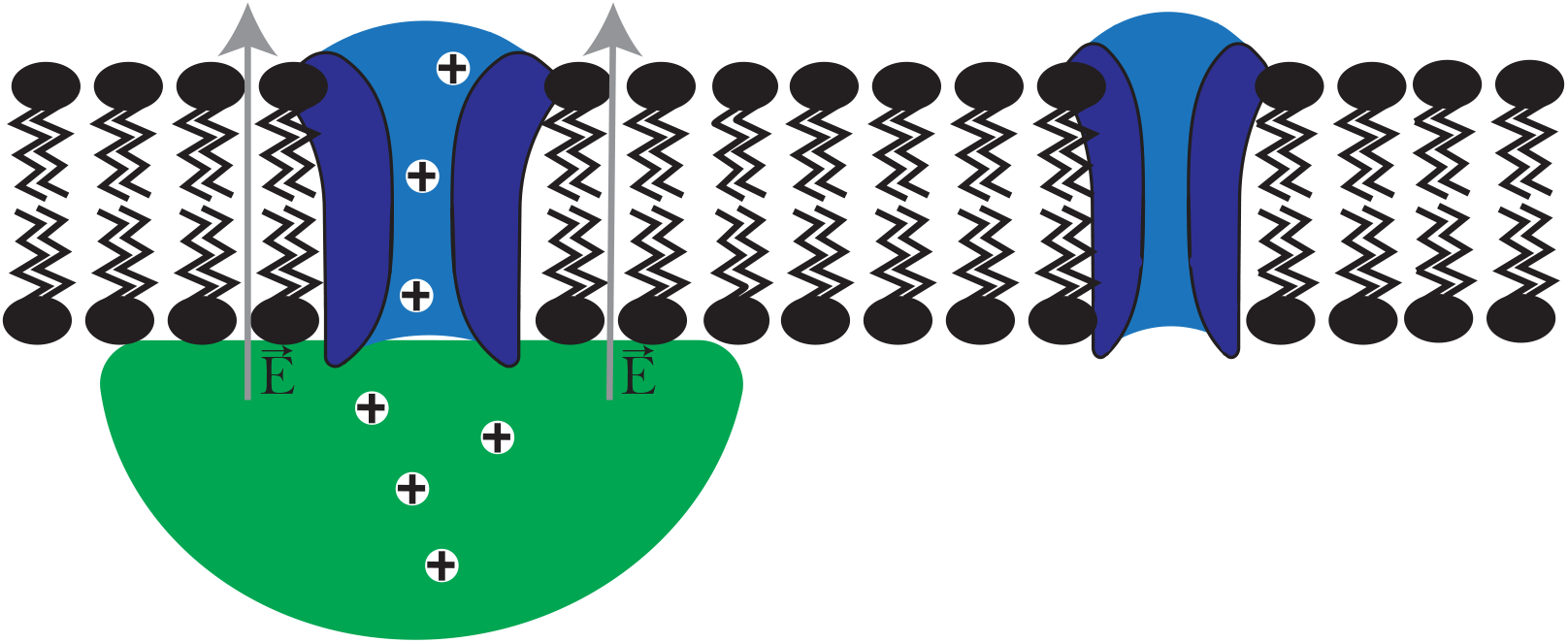
Illustration of hypothetical activation of voltage gated ion channels (blue) by condensates (green) due to induced membrane potential.

Finally, we can also connect our results to two other conceptual directions in the condensate field. In one, recent work has demonstrated the ability of condensate surfaces (and associated potentials) to alter the kinetics of covalent chemical reactions such as the hydrolysis of ATP [50, 51]. We speculate that similar effects might occur within or adjacent to the membrane regions perturbed by interactions with charged condensates, an exciting direction we aim to pursue in future work. In another, we note that while we only explored the role of relatively large micron-scale condensates in this work, our findings should also apply to interactions of nanoscale charged condensates (e.g., percolation clusters [52-54]) with membranes, as also suggested by our simulations. Thus, exploring for induced local membrane potentials and their biological consequences in such a context (for example, in the synaptic region) is another interesting future direction.

## Acknowledgements

The authors gratefully acknowledge support from the National Institutes of Health (NIH/NIGMS Grant R35 GM130375 to A.A.D.). K.L. acknowledges the Gordon & Betty Moore Foundation for support of this work through the Moore Inventor Fellowship number 579361. A.G. gratefully acknowledges support from the Professor Ian A. Wilson Endowed Fellowship for structural biology in the Skaggs Graduate School of Chemical and Biological Sciences. We also thank the various lab members of the Deniz, Lasker and Racki Labs for their thoughtful contributions and discussion. We also thank Leslie Loew for advice regarding potentiometric dyes.

## Materials and Methods

Detailed materials and methods are provided in the supplementary material.

1-palmitoyl-2-oleoyl-sn-glycero-3-phosphocholine (POPC), and 1,2-dioleoyl-sn-glycero-3-phosphate (PA) were purchased from Avanti Polar Lipids (Alabaster, AL). 4-[2-[6-(dioctylamino)-2-naphthalenyl]ethenyl]-1-(3-sulfopropyl)-pyridinium (di-8-anepps) was purchased from Cayman Chemical (Ann Arbor, Michigan). Mineral Oil (USP grade) was purchased from Target (Up & Up). Poly-L-lysine hydrochloride (polyK) was purchased from Alamanda Polymers (Huntsville, AL). ATP disodium salt hydrate (587.17 Da) was purchased from MedChemExpress (Monmouth Junction, NJ). ATTO 565 NHS-ester was purchased from ATTO-TEC (Siegen, Germany).

### Sample Preparation

The Inverted emulsion method was used to generate Giant Unilamellar Vesicles (GUVs) [55]. For imaging, 4 µL of the GUV suspension was added to a coverslip followed by 1 µL of dye solution (10 µM di-8-anepps, 50 mM Tris-HCl (pH 7.5), 500 mM Glucose) and 5 µL of preformed condensates (in 500 mM glucose, 50 mM Tris-HCl (pH 7.5). The preformed condensates were made at 2X concentration and incubated in 0.65 mL tubes for 30 min prior to imaging. A more detailed version of the methods is given in Supplementary Materials.

### Imaging and Analysis

Epifluorescence microscopy was performed on a Nikon Ti2-E automated inverted microscope with a 100x oil immersion objective. For ratiometric images, the dynamic range of the ratios was reduced for visualization purposes by setting cutoff maximum and minimum ratios and assigning values greater or less than those cutoffs to the maximum or minimum ratio respectively.

In ImageJ, polygonal ROIs were drawn corresponding to the membrane segment wetted by the condensate, the free membrane segment, the background within the condensate, and the background adjacent to the free membrane (outside the GUV). The mean intensity for all ROIs were exported Microsoft Excel, and the final background subtracted intensities were calculated as (I_wetted membrane_ - I_condensate background_) and (I_free membrane_ –I_free background_). Using the background subtracted intensities, we calculated the excitation ratios R_440/514_ for the “free membrane” and “wetted membrane” segments. We then calculated the percent change in R_440/514_ using the equation, 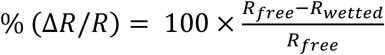, and this quantity was subsequently converted to change in voltage using the equation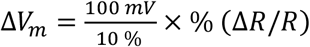. The conversion factor 10%/100 mV has been commonly used in the literature [42, 56].

