## Supplementary Material for "Biomolecular Condensates can Induce Local Membrane Potentials"

(Supplementary Figures, Note-*Analytical Framework* & Materials and Methods)

Anthony Gurunian, Keren Lasker & Ashok A. Deniz*

Department of Integrative Structural and Computational Biology, The Scripps Research Institute,
10550 N. Torrey Pines Rd., La Jolla, CA 92037

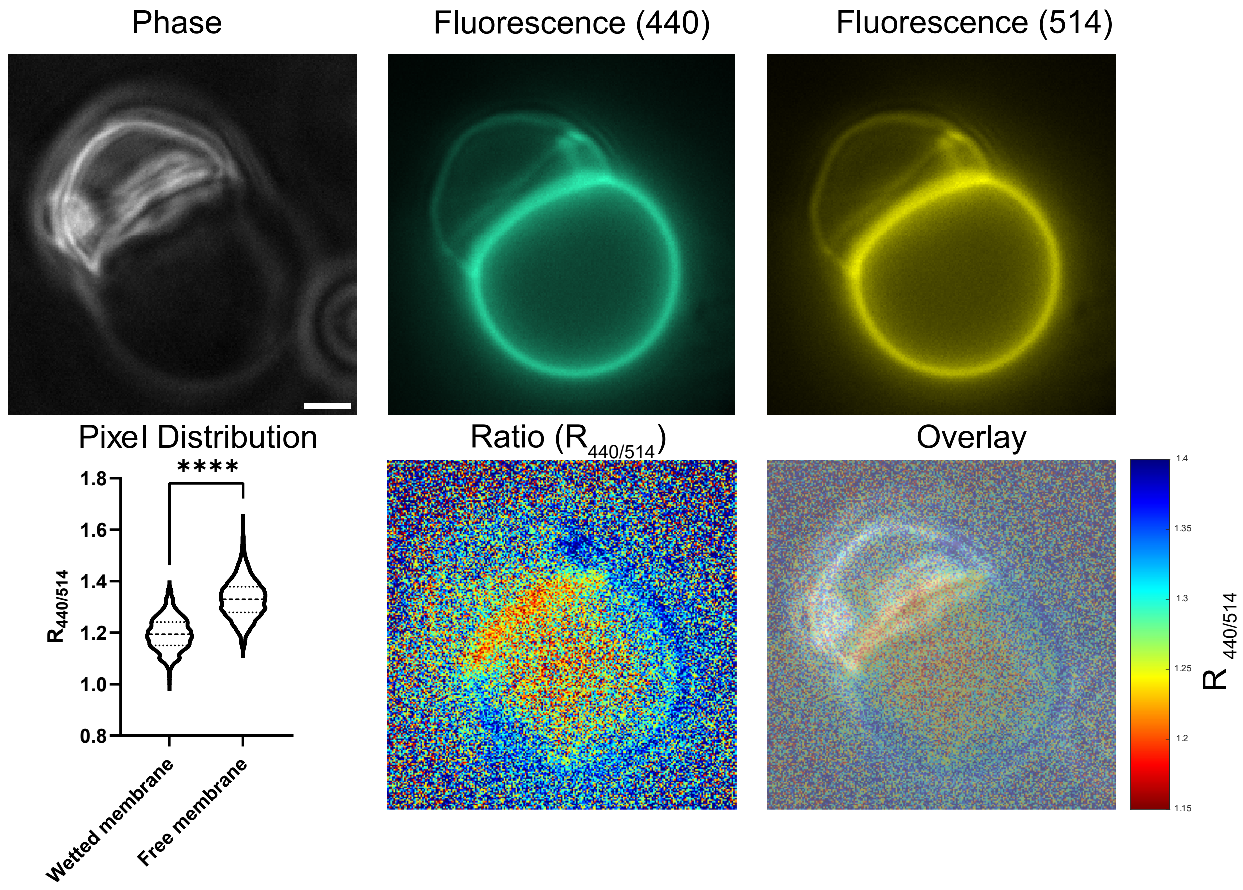


**Figure S1:** Same as main-text Figure 2, with unprocessed raw ratiometric images.

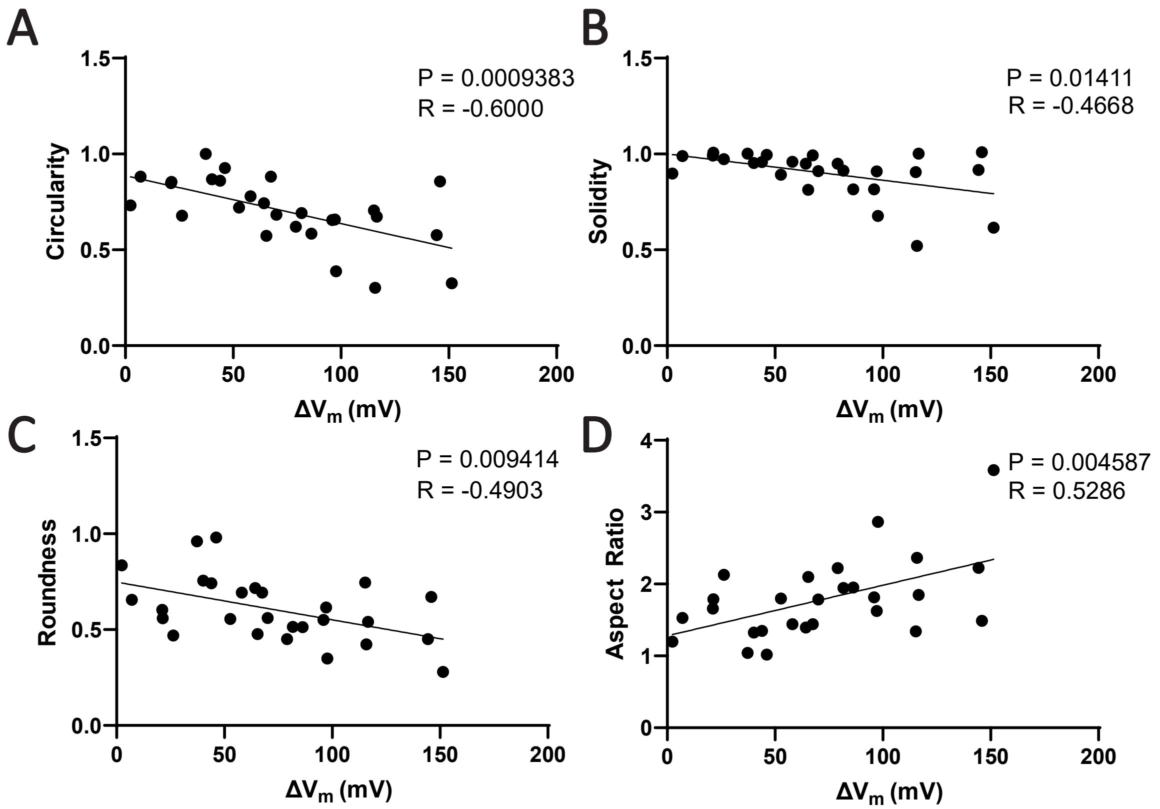


**Figure S2:** Correlation of condensate shape parameters with $\Delta V_m$. Condensates (20 μM polyK-50, 20 μM polyK-100, 1 mM ATP, 2.5 mM $MgCl_2$). R value is Pearson's correlation coefficient. Definitions: Circularity = $\frac{4\pi \times \text{Area}}{perimeter^2}$; Solidity = $\frac{Area}{Convex\ Area}$; Aspect Ratio = $\frac{Major\ Axis}{Minor\ Axis}$; Roundness = $\frac{4 \times \text{Area}}{\pi \times Major\ Axis^2}$

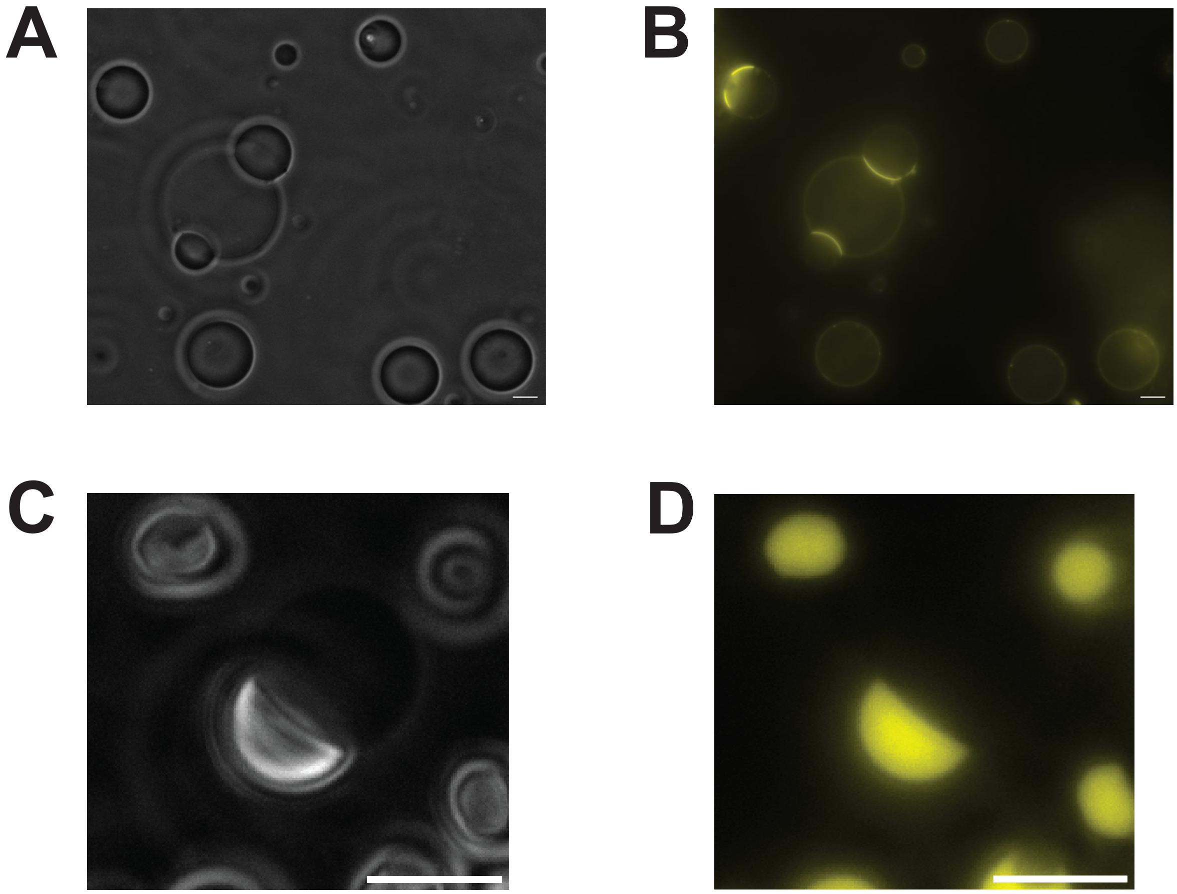


**Figure S3:** Top Row: 20% PA, 80% POPC GUVs with 150 nM polyK-Atto565 only. (A) Phase contrast image (B) Epifluorescence image Bottom row: condensates are formed with 20 μM polyK-50, 20 μM polyK-100, 1 mM ATP, 2.5 mM $MgCl_2$, and 150 nM polyK-Atto565. (C) Phase contrast image (D) Epifluorescence image. Scale bar is 5 μm.

### Supplementary Note 1 - Analytical Framework

Consider a 3-component system consisting of a polycation, anion, and water. Briefly, the electrostatic potential is determined by the distribution of charges via the Poisson equation [1]:

$$\nabla^2\psi = -\frac{\rho(\boldsymbol{r})}{\varepsilon} \tag{1}$$

where $\rho(\boldsymbol{r}) = \sum_{i=1}^{3} z_i e n_i(\boldsymbol{r})$ is the charge density. The free-energy functional is given by:

$$F[n_i, \psi] = \int f_{FH}(n_i) + \frac{\kappa_i}{2}\left(\nabla n_i(\boldsymbol{r})\right)^2 + \rho(\boldsymbol{r})\psi(\boldsymbol{r}) \tag{2}$$

where $n_i$ is the local concentration of component $i$, $\psi$ is the electric potential, $\kappa_i$ is the gradient cost for component $i$, and $f_{FH}(n_i)$ is the Flory-Huggins free energy given by:

$$\frac{f_{FH}(n_i)}{k_B T} = \sum_{i=1}^{3} n_i \, ln \, v_i n_i(\boldsymbol{r}) + \sum_{i<j} \chi_{ij} n_i(\boldsymbol{r}) n_j(\boldsymbol{r}) \tag{3}$$

where $v_i$ is the molecular volume of component $i$, and $\chi_{ij}$ is the Flory-Huggins interaction parameter between components $i$ and $j$. The electrochemical potential is then given by:

$$\mu_i(\boldsymbol{r}) = \frac{\delta F}{\delta n_i} = \frac{\partial F}{\partial n_i} - \nabla \cdot \frac{\partial F}{\partial \nabla n_i} = \frac{\partial f_{FH}}{\partial n_i} - \kappa \nabla^2 n_i(\boldsymbol{r}) + z_i e \psi(\boldsymbol{r}) \tag{4}$$

where

$$\frac{\partial f_{FH}}{\partial n_1} = k_B T \left[ ln\left(v_1 n_1(\boldsymbol{r})\right) + 1 + \frac{v_1}{v_3}\left(1 - ln\left(v_1 n_1(\boldsymbol{r})\right)\right) + \chi_{12} n_2(\boldsymbol{r}) + \chi_{13} n_3(\boldsymbol{r}) - \frac{v_1}{v_3}\left(\chi_{13} n_1(\boldsymbol{r}) + \chi_{23} n_2(\boldsymbol{r})\right) \right] \tag{5}$$

$$\frac{\partial f_{FH}}{\partial n_2} = k_B T \left[ ln\left(v_2 n_2(\boldsymbol{r})\right) + 1 + \frac{v_2}{v_3}\left(1 - ln\left(v_2 n_2(\boldsymbol{r})\right)\right) + \chi_{12} n_1(\boldsymbol{r}) + \chi_{23} n_3(\boldsymbol{r}) - \frac{v_2}{v_3}\left(\chi_{13} n_1(\boldsymbol{r}) + \chi_{23} n_2(\boldsymbol{r})\right) \right] \tag{6}$$

Note that the concentration of component 3 is determined by conservation of volume fraction:

$$v_3 n_3(\boldsymbol{r}) = 1 - v_1 n_1(\boldsymbol{r}) - v_2 n_2(\boldsymbol{r}) \tag{7}$$

To determine the equilibrium concentration profiles, and equilibrium electric potential profile we solve the following equation [2]:

$$\frac{dn_i(\boldsymbol{r})}{dt} = -\mu_i(\boldsymbol{r}) + \langle \mu_i(\boldsymbol{r}) \rangle \tag{8}$$

with the boundary conditions:

$$\left.\frac{dn_i}{dx}\right|_{x=0,N} = 0; \left.\frac{d\psi}{dx}\right|_{x=0} = -\frac{\sigma}{\varepsilon}; \left.\frac{d\psi}{dx}\right|_{x=N} = 0 \tag{9}$$

which corresponds to a charged boundary with surface charge density $\sigma$ at $x = 0$. The surface charge density is determined by

$$\sigma = \frac{e}{A_L} f[\theta(z-1) - (1-\theta)] \tag{11}$$

where $e$ is the elementary charge, $A_L$ is the area per lipid, $f$ is the fraction of charged lipids, and $\theta$ is the fractional occupancy of binding sites which is determined by the Langmuir Isotherm:

$$\theta = \frac{K_{p+} n_{p+}\big|_{x=0,N}}{1 + K_{p+} n_{p+}\big|_{x=0,N}} \tag{12}$$

We solve equation 9 with the appropriate boundary conditions (eq. 10) by discretizing the spatial derivatives using central finite differences, and integrate the time derivatives using MATLAB's ode15s() function. All code used in this paper will be made available at (github.com/Deniz-Lab)

### Supplementary Materials and Methods

1-palmitoyl-2-oleoyl-sn-glycero-3-phosphocholine (POPC), and 1,2-dioleoyl-sn-glycero-3-phosphate (PA) were purchased from Avanti Polar Lipids (Alabaster, AL). 4-[2-[6-(dioctylamino)-2-naphthalenyl]ethenyl]-1-(3-sulfopropyl)-pyridinium (di-8-anepps) was purchased from Cayman Chemical (Ann Arbor, Michigan). Mineral Oil (USP grade) was purchased from Target (Up & Up). Poly-L-lysine hydrochloride (polyK) was purchased from Alamanda Polymers (Huntsville, AL). ATP disodium salt hydrate (587.17 Da) was purchased from MedChemExpress (Monmouth Junction, NJ). ATTO 565 NHS-ester was purchased from ATTO-TEC (Siegen, Germany).

*Sample Preparation*

The Inverted emulsion method was used to generate Giant Unilamellar Vesicles (GUVs) [3]. Briefly, lipids from stock solutions in chloroform (10 mg/mL) were dissolved in mineral oil for a final concentration of 0.6 mg/mL and heated at 80°C in a fume hood to evaporate the chloroform.

First, 200 µL outer aqueous solution (50 mM Tris-HCl (pH 7.5), 500 mM Glucose) was added to a 1.5 mL microcentrifuge tube. In a separate 2 mL microcentrifuge tube, 50 µL inner aqueous solution (50 mM Tris-HCl (pH 7.5), 500 mM Sucrose) was combined with 400 µL of lipids in oil (0.15 mg/mL lipid) and vortexed on high for 40-60 seconds to create an emulsion. The emulsion was then gently pipetted on top of the bottom aqueous solution and incubated at room temperature for 20 minutes to equilibrate the oil-water interface. The tube was then centrifuged at 16000 g for 15 minutes, producing a pellet of vesicles. The oil layer and part of the aqueous layer was pipetted off, and the pellet were resuspended in 100 µL of bottom aqueous solution. The 100 µL vesicle suspension was then combined with 400 µL of bottom aqueous solution in a new 1.5 mL microcentrifuge tube and centrifuged at 5000 g for 5 minutes. Part of the bottom aqueous solution was removed, and the pellet was resuspended in a final volume of 100-200 µL. For imaging, 4 µL of the GUV suspension was added to a coverslip followed by 1 µL of dye solution (10 µM di-8-anepps, 50 mM Tris-HCl (pH 7.5), 500 mM Glucose) and 5 µL of preformed condensates (in 500 mM glucose, 50 mM Tris-HCl (pH 7.5). The preformed condensates were made at 2X concentration and incubated in 0.65 mL tubes for 30 min prior to imaging. Coverslips were passivated with 10-20 mg/mL BSA for at least 24 hours and rinsed with DI water prior to imaging.

We noticed that the ATP concentrations of liquid stocks made by weighing out ATP were consistently 15% lower than the ATP concentrations determined by measuring the absorbance at 260 nm on a NanoDrop2000c, using an extinction coefficient of 15,400 $cm^{-1}$ $M^{-1}$. We corrected our ATP concentrations to match the concentration determined using the absorbance.

*Imaging and Analysis*

Epifluorescence microscopy was performed on a Nikon Ti2-E automated inverted microscope with a 100x oil immersion objective (Plan Apo λ 100x Oil Ph3 DM) and a Kinetix sCMOS camera (Teledyne Photometrics). Images were acquired with the NIS Elements software version

5.42.03. To generate ratiometric pixel distributions, ROIs were drawn in MATLAB, and the list of individual pixel intensity ratios for each ROI was exported to GraphPad Prism for graphing and statistics. For ratiometric images, the dynamic range of the ratios was reduced for visualization purposes by setting cutoff maximum and minimum ratios and assigning values greater or less than those cutoffs to the maximum or minimum ratio respectively. For the ratiometric image in Figure 2, the noisy background was set to a constant (background) value by using local intensity thresholds determined by adaptive window sizes based on the local intensity gradient. All code used in this paper will be made available at (github.com/Deniz-Lab). Phase contrast images were taken with 100 ms exposure and a Nikon DAPI 96360 filter cube. Epifluorescence images were taken using 440 nm and 514 nm excitation with 20-30 ms exposure and a Nikon Spectra-Triple filter cube (Emission 464 nm - 486 nm, 532 nm - 554 nm, 603 nm – 800 nm). Image resolution was 15.14 pixels per micron.

To generate summary results/figures, the following disqualifying criteria were used to curate data prior to processing: 1. No clear membrane segment between condensate and membrane; 2. Signal to background ratio less than 3 in the 440 nm channel; 3. Condensate moved between frames; 4. GUV or condensate out of focus; 5. Background artifacts near vesicle; 6. GUV diameter < 2µm; 7. Pixel Saturation; 8. Complete wetting of GUV (no free membrane segment) 9. GUV cut off at edge of field of view. 10. Other uninterpretable features. When measuring ROIs we also followed two additional rules. 1. If more than one condensate or GUV is contacting the GUV of interest, measure the union of the free membrane segments as the "free membrane". 2. If another vesicle is adsorbed to the GUV of interest, do not measure the membrane segment at the interface of the two vesicles.

In ImageJ, polygonal ROIs were drawn corresponding to the membrane segment wetted by the condensate, the free membrane segment, the background within the condensate, and the background adjacent to the free membrane (outside the GUV). The mean intensity for all ROIs were exported to Microsoft Excel, and the final background subtracted intensities were calculated as ($I_{\text{wetted membrane}}$ - $I_{\text{condensate background}}$) and ($I_{\text{free membrane}}$ –$I_{\text{free background}}$). Using the background subtracted intensities, we calculated the excitation ratios $R_{440/514}$ for the "free membrane" and "wetted membrane" segments. We then calculated the percent change in $R_{440/514}$ using the equation, $\%\ (\Delta R/R) = 100 \times \frac{R_{free} - R_{wetted}}{R_{free}}$, and this quantity was subsequently converted to change in voltage using the equation $\Delta V_m = \frac{100\ mV}{10\ \%} \times \%\ (\Delta R/R)$. The conversion factor 10%/100 mV has been commonly used in the literature [4, 5].

*FRAP*

For FRAP (Fluorescence recovery after photobleaching) experiments, 300 nM polyK100-Atto565 was doped into the polyK/ATP coacervates. The bleach radius was set to 0.2 µm in the center of the droplet, The stimulation dwell time was set to 100 µs, and stimulation was done with 405 nm (20% intensity), and 561 nm (40% intensity) lasers. Acquisition was at 561 nm with 50 ms exposure. Bleached droplets were around 4-8 microns in diameter. FRAP curves were calculated using the equation $I_N(t) = \frac{I(t) - I_0}{R(t) - I_0}$ where $I_N$ is the normalized intensity, $I(t)$ is the raw intensity of the bleached ROI at time $t$, $I_0$ is the raw intensity immediately after bleaching, and $R(t)$ is the raw intensity of a reference droplet at time $t$.

*polyK labeling*

We followed a labeling protocol from Thermo Scientific for NHS-Fluorescein (https://assets.thermofisher.com/TFS-Assets/LSG/manuals/MAN0011647_NHSFluorescein_UG.pdf) and adapted it for labeling polyK-100 with ATTO 565 with minor modifications. Briefly, we incubated polyK-100 with ATTO 565 NHS-ester at a 1:1 molar ratio for 1 hour in the dark at room temperature in a 20 mM HEPES (pH 8.2) buffer. We then removed excess dye by passing the sample through a Pierce™ Dye Removal Column (ThermoFisher catalog no. A44296). Next, we filtered the sample with an Amicon® Ultra-0.5 Centrifugal Filter (3000 MWCO) to buffer exchange into Milli-Q water. The final stock concentration of polyK-Atto565 was approximately 30 μM, and was stored in the -20 Freezer for future use.
